# Combined direct/indirect detection allows identification of DNA termini in diverse sequencing datasets and supports a multiple-initiation-site model for HIV plus-strand synthesis

**DOI:** 10.1101/2023.06.12.544617

**Authors:** William Wang, Karen L. Artiles, Shinichi Machida, Monsef Benkirane, Nimit Jain, Andrew Z. Fire

## Abstract

Replication of genetic material involves the creation of characteristic termini. Determining these termini is important to refine our understanding of the mechanisms involved in maintaining the genomes of cellular organisms and viruses. Here we describe a computational approach combining direct and indirect readouts to detect termini from next-generation short-read sequencing. While a direct inference of termini can come from mapping the most prominent start positions of captured DNA fragments, this approach is insufficient in cases where the DNA termini are not captured, whether for biological or technical reasons. Thus, a complementary (indirect) approach to terminus detection can be applied, taking advantage of the imbalance in coverage between forward and reverse sequence reads near termini. A resulting metric (“strand bias”) can be used to detect termini even where termini are naturally blocked from capture or ends are not captured during library preparation (e.g., in tagmentation-based protocols). Applying this analysis to datasets where known DNA termini are present, such as from linear double-stranded viral genomes, yielded distinct strand bias signals corresponding to these termini. To evaluate the potential to analyze a more complex situation, we applied the analysis to examine DNA termini present early after HIV infection in a cell culture model. We observed both the known termini expected based on standard models of HIV reverse transcription (the U5-right-end and U3-left-end termini) as well as a signal corresponding to a previously described additional initiation site for plus-strand synthesis (cPPT [central polypurine tract]). Interestingly, we also detected putative terminus signals at additional sites. The strongest of these are a set that share several characteristics with the previously characterized plus-strand initiation sites (the cPPT and 3’ PPT [polypurine tract] sites): (i) an observed spike in directly captured cDNA ends, an indirect terminus signal evident in localized strand bias, (iii) a preference for location on the plus-strand, (iv) an upstream purine-rich motif, and (v) a decrease in terminus signal at late time points after infection. These characteristics are consistent in duplicate samples in two different genotypes (wild type and integrase-lacking HIV). The observation of distinct internal termini associated with multiple purine-rich regions raises a possibility that multiple internal initiations of plus-strand synthesis might contribute to HIV replication.

## Introduction

Extrachromosomal DNA, which occurs in both circular and linear forms, plays a significant role in numerous biological processes such as gene amplification in cancer^1,2^ and microbial biofilm formation.^3,4^ Although technologies have been devised to study circular extrachromosomal DNA^5^, investigations of linear extrachromosomal DNA elements have been more limited in part because of the difficulty in distinguishing integrated from extrachromosomal molecules.^6–8^ Linear extrachromosomal DNA has been found in biological systems^9^, including linear plasmids in *Actinoplanes*^10^, Lyme disease isolates^11,12^, and the mitochondria of many plant and fungi species^13,14^. Linear extrachromosomal ribosomal DNA plasmids also exist in *Candida albicans*.^15^ Similarly, reverse-transcription in cells infected with different retroviruses also produces linear extrachromosomal DNA.^16^ To understand their pertinent properties and functions, tracking linear extrachromosomal DNA is necessary, and identification of linear extrachromosomal DNA (compared to circular or chromosomally integrated forms) is essential.

In contrast to circular DNA, linear DNA is definitionally distinguished as having termini or ends. Previous methodologies for detecting DNA termini have leveraged a large number of experimental tools^17^, with experimental detection remaining the gold standard for definitive assignment of topological characteristics. However, because experimental techniques to measure topology and infer termini require time and resources that may be limiting, and because most of the publicly available next-generation sequencing data have been acquired using standard protocols that do not explicitly aim to determine termini, there is interest in computational approaches to infer potential termini from the vast majority of sequencing data that are already available.

Previously published computational approaches to identify termini from next-generation sequencing data have generally relied on an increased likelihood of capture for consistent termini that are present in a biological sample.^18–20^ The idea here relies on fragmentation of DNA for sequencing resulting in a profile of captured ends that randomly samples nucleotides in the genome. Overlaying on this pattern, a discontinuity resulting from a consistent break at a specific position that can also be captured for sequencing will produce a strong peak in the observed profile of read termini. This direct method effectively detects ends in experimental data, provided three criteria are met:

1. Sequencing protocols entail fragmentation of input DNA followed by capture of ends.
2. Termini pre-existing in the pre-processed DNA are preferentially captured.
3. Fragmentation and capture are sufficiently random (and termini sufficiently prevalent) that true ends will show a positive enrichment signal when compared to neighboring sequences.

Methods based on the above have been useful for determining bacteriophage termini and terminal sequences in randomly fragmented sequencing data and metagenomic virome datasets, for example.^18–20^ But in some cases, the nature of the sequencing protocols or ends precludes direct application of such an algorithm. In particular, DNA termini that are not available for capture, or methods that only capture internal DNA fragments within a molecule (e.g., tagmentation-based methods such as “Nextera”) may result in an inability to observe true termini.

In this work, we use an additional approach that exploits an independent signature of DNA termini in sequencing datasets: sequencing strand bias (also referred to in other work as unbalanced strand mapping^21^). While previously noted in a number of reports as a characteristic for detection of RNA ends and DNA replication forks,^22,23^ we are unaware of any current tools that apply this property to the general identification of termini. Here, we describe and apply the detection of such strand bias as a general approach to end discovery, with specific application to understanding the termini present in data from sequencing of viral samples in the context of a complex animal genome. The approach we describe is particularly suited for identification of linear extrachromosomal DNA such as viruses with well-defined ends.

## Results

### Model for strand bias and end capture

The termini detection method outlined here is evident from a careful consideration of a hypothetical sequencing experiment in which a pre-existing linear template DNA is experimentally fragmented into “shotgun fragments” that are captured and sequenced from one or both ends (Fig. 1a and 1c). We consider two types of reads near pre-existing template termini. Outward-facing reads, although contained within the fragment of interest, are sufficiently close to a terminus in the original template that the underlying fragment (generally longer than the actual read) would press right up against (or overlap) the defined terminus. Inward-facing reads are those whose position presents little or no constraint on extension. Regardless of the underlying sequencing technology, coverage from inward-facing reads will be observed continuously from the pre-existing terminus, while outward-facing reads will be lost for any area close to such a terminus (Fig. 1). If shotgun fragments have a uniform length L, then outward-facing reads can only start at least L bases from a terminus. In cases where fragment length is somewhat variable, reads adjacent to pre-existing termini will still be highly enriched in the inward-facing direction. A positional map of relative sequence representation using coverage on the two strands (sense and antisense reads), which we will refer to as a strand bias plot, can thus provide us with a provisional picture that identifies candidate termini (Fig. 1b and 1d).

**Fig. 1.**
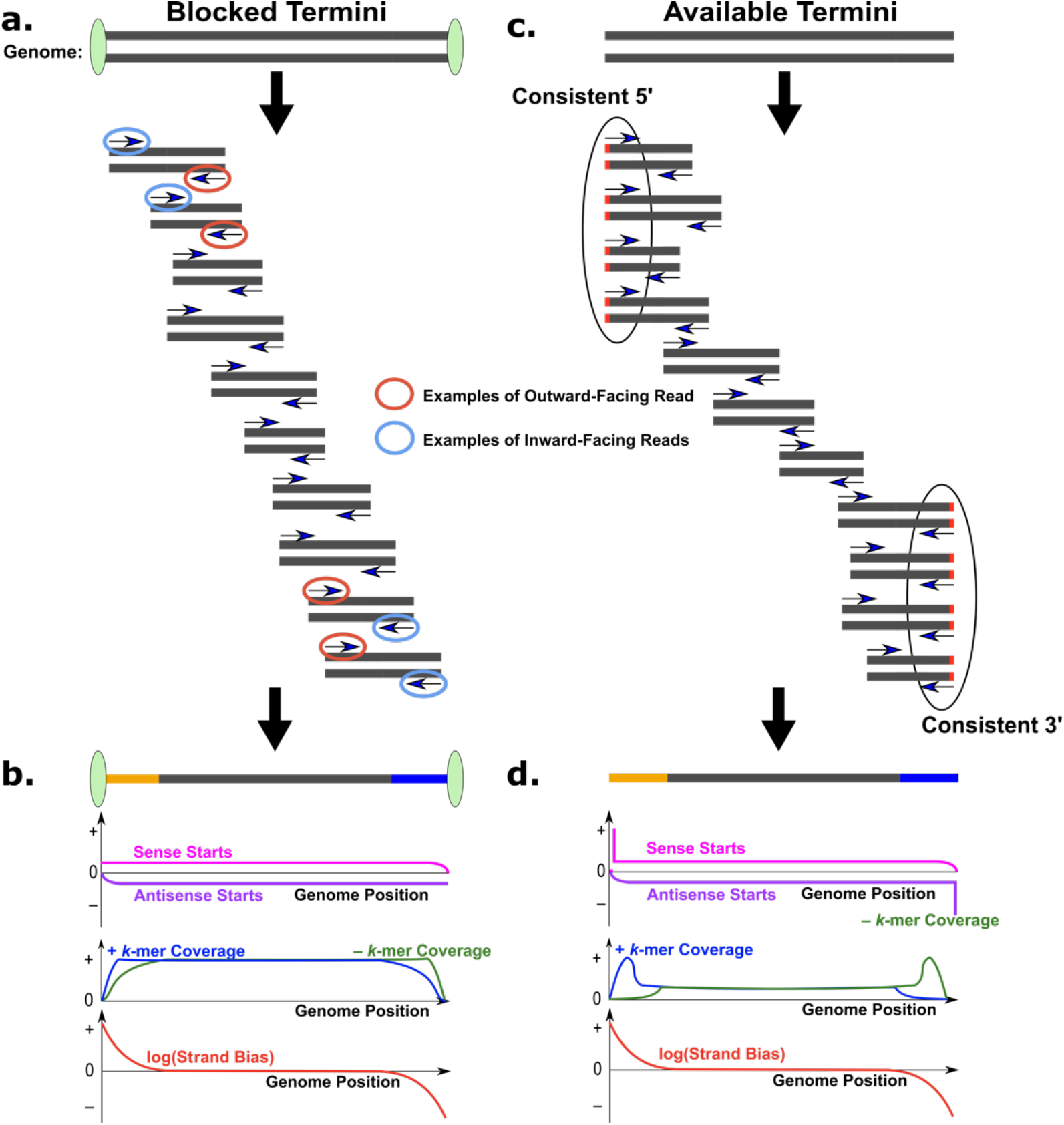
Next-generation sequencing schematic and impact on end representation. Fragmentation and sequencing of a genome with **a**, blocked termini and **c**, available termini. Resulting end capture, *k*-mer coverage, and strand bias for the case of **b**, blocked ends and **d**, available ends.^24^ **b and d**, Genome depicted with termini regions highlighted in gold and blue. Plots show coverage of sense (pink) and antisense (purple) DNA sequencing read starts, *k*-mer coverage for plus (blue) and minus (green) strand and calculated strand bias (red). **d**, Sequencing reads (blue arrows) captured from unblocked ends yield consistent 5’ and 3’ ends (black ovals) that are enriched in coverage. Here, strand bias is defined as + Coverage/– Coverage.

If the fragmentation is close-to-random and there are no existing termini (e.g., a circular template), we expect equal amounts of sequence representation in the plus (5’-3’, +) and minus (3’-5’, ?) strands, which holds for regions of linear DNA that are not near defined termini.

Sequences near a terminus will, by contrast, exhibit biased representation (provided that fragment length, is greater than read length); that is, the representation of plus and minus strands is unequal (Fig. 1b and 1d). Additionally, for protocols that fragment input DNA and capture ends, a desired broadly distributed fragmentation during library preparation will yield datasets in which true DNA termini are preferentially represented (Fig. 1c and 1d). This leads to much higher sense and antisense start positions at DNA termini when there are accessible ends (Fig. 1d).

We use several different sets of tools to visualize and understand strand-specific representation of individual subregions in next-generation sequencing datasets. In one approach, we first align the sequencing dataset to a matching reference genome using Bowtie 2 or BWA.^25-^ 27 Coverage is then calculated as a function of position in a reference genome for each strand, and the relative coverage is plotted to identify regions where the two strands are represented differently.

To assess strand bias using an alignment-independent approach, we also describe our tool for the counting of individual nucleotide strings (*k*-mers) in sense and antisense orientations and comparison of these counts. Although still subject to a variety of concerns and caveats, the *k*-mer based approaches provide an orthogonal means for coverage quantitation that can help avoid the pitfalls of alignment-based read counting. Metrics based on *k*-mer counting use a variety of relatively simple tools that we have adapted for termini identification. Our counting and display script, named PolyBench, computes coverage, strand bias, and start/end positions, which are depicted on the same plot, allowing for convenient observation of DNA termini via a number of methods. PolyBench is available on our GitHub repository at https://github.com/FireLabSoftware/PolyBench.

### Initial proof-of-concept: Analysis of bacteriophage lambda DNA

To test our strand bias metric, we created a defined set of linear molecules by digesting *Escherichia coli* phage lambda genomic DNA^28^ with a restriction enzyme (we used either *Pst*I, which generates sticky ends with a 4 base 3’ overhang or *Pvu*II, which generates blunt ends^29,30^) and sequenced that alongside intact (uncut/control) DNA. Sequencing was performed using a tagmentase-based DNA fragmentation method (Nextera). We aligned DNA sequencing reads to an established lambda reference genome (see Supplementary Material) and plotted strand bias, which is defined as

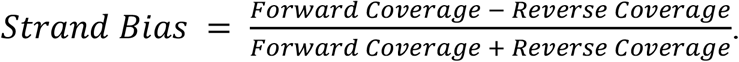

We observe, as predicted by our model, a strand bias peak at each location where the enzyme cuts the DNA,^31^ as demonstrated by the overlap of the peaks with the triangle markers representing enzyme cut sites (Fig. 2). Although the magnitude of the peaks clearly distinguishes the true signal (of enzyme-digested DNA termini) from the noise (resulting from fragmentation pattern biases of tagmentase), subtracting the control (uncut) datasets demonstrated a clearer isolation of the true signal peaks. (Fig. 2). By subtracting the control condition from the test condition (digested DNA), we can extract only the signal from the linear DNA while minimizing systematic fragmentation biases from the sequencing method.

**Fig. 2.**
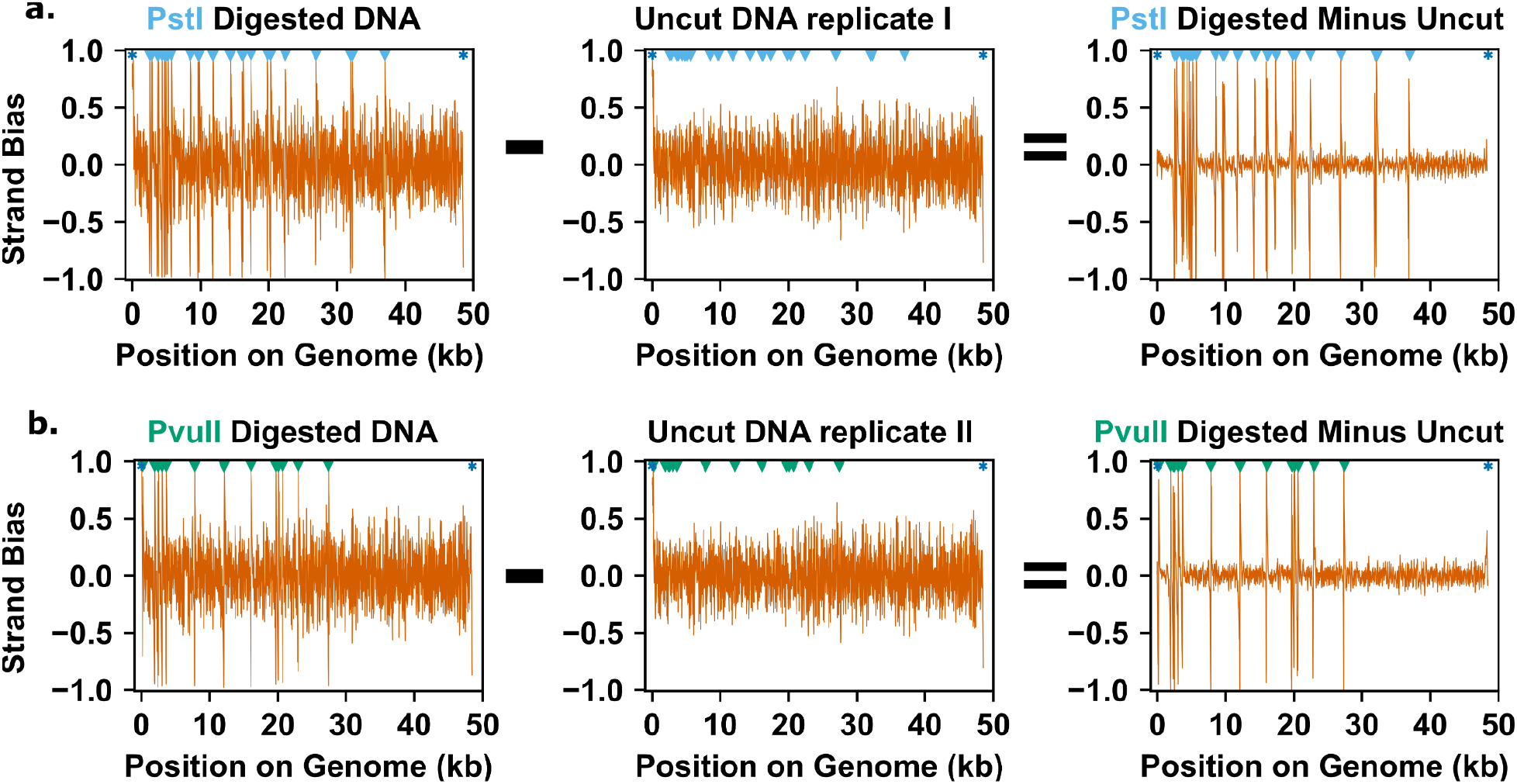
Test of the strand bias model using restriction enzyme digestion of bacteriophage lambda DNA. Strand bias was plotted for either **a**, PstI-digested, or **b**, PvuII-digested DNA along with a corresponding uncut DNA control. Plots showing the remaining signal after subtracting the uncut signal from the restriction enzyme-treated signal are shown in the right panels. For this analysis, we have enforced a minimum size requirement on shotgun fragments of at least 75 base pairs, which is less than the read length (78), with alignment using BWA^17^. Triangles correspond to enzyme cut sites and asterisks correspond to ends of the genome. Note that no threshold on coverage was used.

### Detection of linear DNA during adenovirus infection

Certain mammalian viruses (e.g., adenoviruses) have linear double-stranded DNA (dsDNA) genomes.^32–36^ We used a publicly available dataset of shotgun DNA sequence reads prepared using Illumina TruSeq following infection by an adenovirus (bovine adenovirus type 7)^37^ to assess the ability of the different terminus detection approaches to illuminate the end structure in a case where ends are blocked. Adenoviruses replicate through a machinery that utilizes a terminal protein primer, resulting in a protein-modified terminus that is not available for ligation;^38^ this mechanism is not unique to adenovirus, but is indeed shared by a number of phage and viruses of eukaryotic cells, including phage phi29, poliovirus, and tectivirus.^39–41^Analysis of the bovine adenovirus dataset to its reference genome exhibited strand bias asymmetries at the start and end of the genome (PolyBench analysis shown in Fig. 3). We interpret this as evidence that bovine adenovirus exists in a linear form, consistent with previous experimental observations.^32–34^ Two significant strand bias peaks are observed: a positive peak at the extreme left of the genome and a negative peak at the far right. In the rest of the genome from 1,000-29,000 base pairs (bp), there are no significant peaks, with only minor peaks, likely occurring from biases in fragmentation. Meanwhile, end capture peaks (represented by “+ Termini-R1”, “– Termini-R1”, “+ Termini-R2”, “– Termini-R2”) are small and scarce, which is expected because the ends are blocked by a protein primer.^38^ As such, the case of adenovirus exemplifies the utility of having both direct and indirect methods available.

**Fig. 3.**
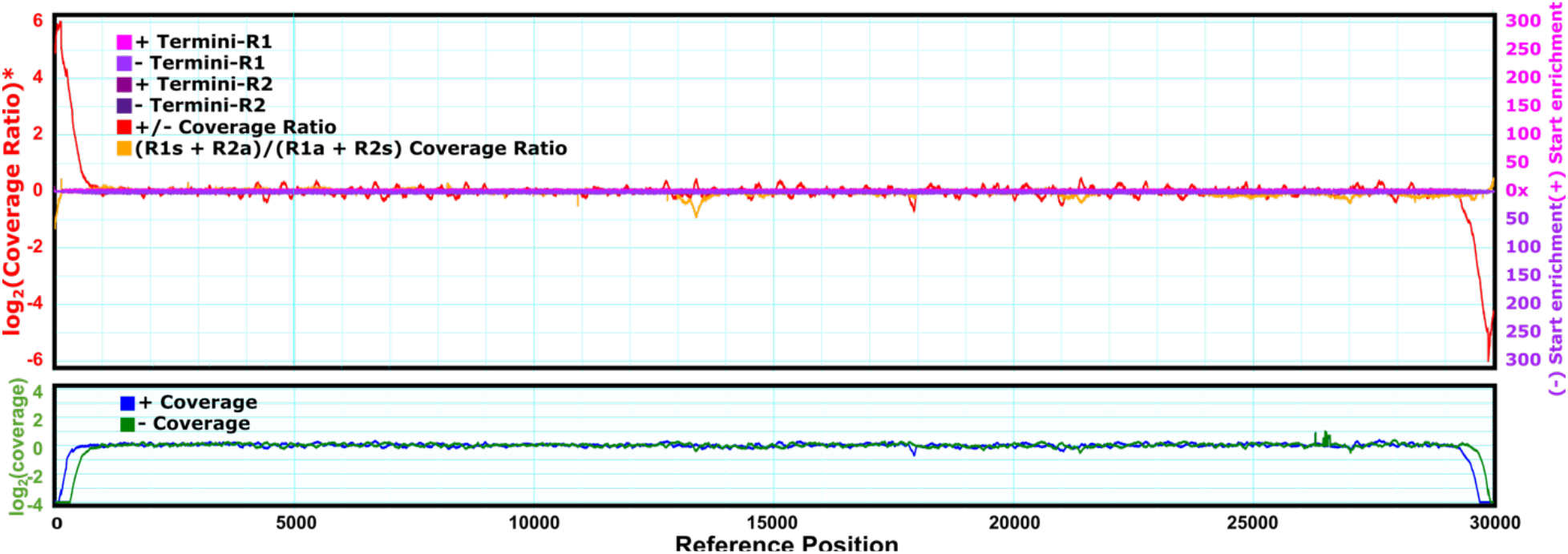
Analysis of sequence data for the blocked linear genome of bovine adenovirus^34^, with display of termini, coverage, and strand bias analysis using PolyBench. Lower plot shows coverage levels for k-mer sequences from read data mapped to the plus-strand (blue) and minus strand (green), with a characteristic shift near the terminus due to the scarcity of outward-facing reads at each end. Top plot shows additional features displayed by PolyBench. Most striking in this case are the dramatic peaks in +/-coverage ratio (ratio of positive to negative strand k-mer counts from PolyBench; red) at the two ends corresponding to the blocked termini. No sites of singular terminus enrichment were observed in this dataset, thus the plotting of single-point terminus enrichment values (“+ Termini-R1”, “-Termini-R1”, “+ Termini-R2”, and “-Termini-R2” all remain near baseline). Cross ratios (R1s+R2a)/(R2a+R1s) are also near baseline. These plots were generated with a *k*-mer length of 27, and ratios have been regressed toward unity through a Bayesian regression (see methods).

### Analysis of DNA termini during early stages of HIV replication

To understand the formation of termini during a more complex viral life cycle, we next analyzed data obtained following infection of human cells with HIV virus. HIV is a retrovirus that introduces RNA into cells along with an active enzyme (reverse transcriptase) capable of converting RNA information into DNA.^42^ A complex set of genome copying steps following infection eventually results in dsDNA molecules with direct repeats at the termini (called long terminal repeats or LTRs). These dsDNA molecules are then integrated into the host genome as part of the viral life cycle. Despite extensive analysis of the intermediates in HIV’s genomic life cycle, there remain several outstanding questions related to this process.^43^

A series of DNA sequencing datasets following HIV infection were available from previous work directed toward understanding of chromatin formation following HIV infection.^35^ That work included a time series following infection [with sequencing performed at 9 hours and 48 hours post infection (hpi)], as well as an option to examine requirements for HIV integrase [enabled by the availability of data for both wild type and integrase mutant HIV with an inactivating mutation at amino acid D116 (IN^D116N^; see methods].^35^ To extend this analysis, we also provide additional data from a 48-hour time point following infection by the integrase mutant. In the HIV datasets used, cellular DNA was processed by sonication to a size of 100-300 bp and analyzed following sequence specific capture of HIV sequences.^35^ For our analysis of HIV data we mapped k-mers and reads using a custom assembled reference genome generated with the assistance of MEGAHIT (see Supplementary Material).^44^

Using our approaches to detect DNA termini, we noted several termini that are enriched at the 9 hpi compared to the 48 hpi. Such differentially enriched termini could represent distinct linear molecular forms that are part of the HIV genome replication cycle or could stem from byproduct replication intermediates. Consistent with a capability to detect linear molecular forms with relevance to HIV replication, we observed high strand bias and end capture peaks at the LTR termini, as expected based on the known HIV replication life cycle (Fig. 4 and Table 1).^45^ As both end capture and strand bias locations are consistent and substantiate the literature, we are confident these represent true ends. Overall, these observations confirm the ability of sequencing asymmetry analysis to detect DNA termini in a complex sample. These results were confirmed in analysis of parallel biological replicate datasets (Fig. S1, Supplementary Material).

**Table 1.**
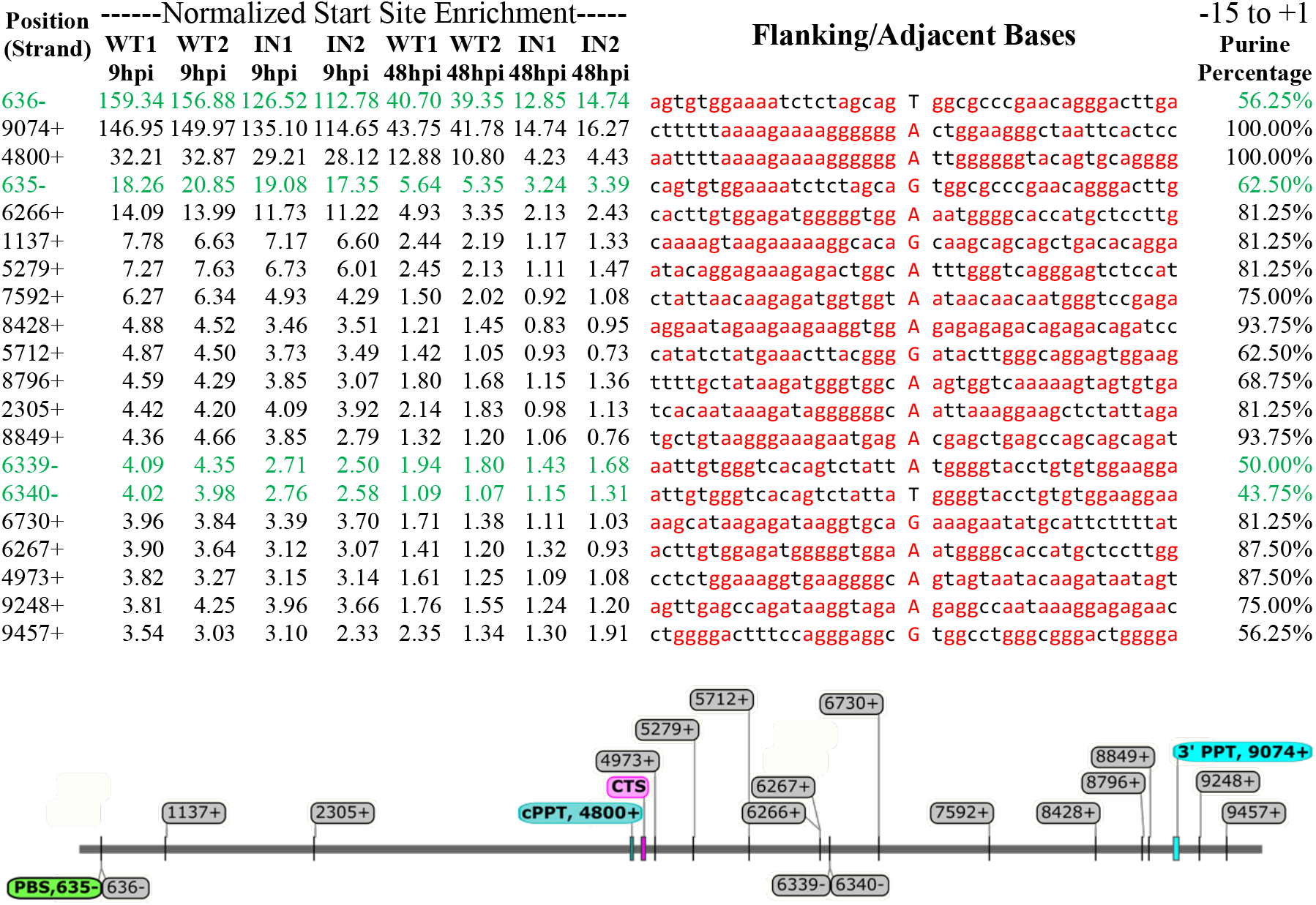
The top 20 start-enriched sites in HIV by end capture (ranked by 9 hpi wild type replicate 1, which is called WT1), including notable known peaks as well as novel purine-rich sequences on the sense strand (altPPTs). Normalized Start Site Enrichment represents R1+R2 divided by TotalStarts/TotalPositions where TotalStarts is the sum of R1s, R2s, R1a, and R2a for all positions. Data for WT1 and IN1 samples correspond to Fig. 4, while the second replicates WT2 and IN2 correspond to Fig. S1 (Supplementary Material). Purine percentages were calculated for 16 nucleotide stretches, corresponding to the known length of the polypurine tracts in HIV. Display of the top 20 sites is arbitrarily presented here; there are also additional sites with weaker start site enrichment and weaker purine bias, with the relative contributions of the individual start sites (shown and not shown) remaining to be experimentally determined.

**Fig. 4.**
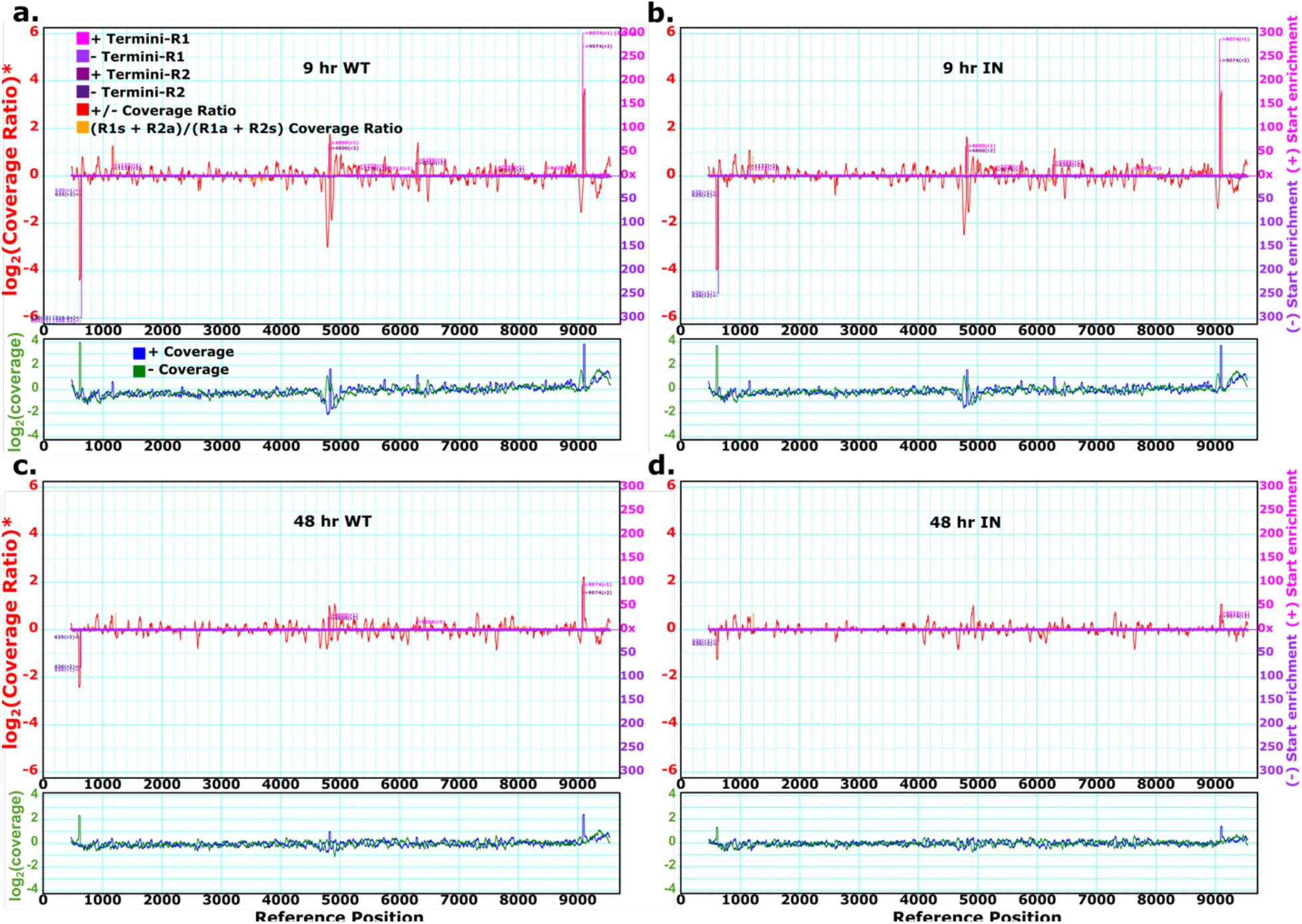
Strand bias and end capture of the HIV genome, calculated by PolyBench, demonstrate DNA termini and [Putative] Plus-Strand Initiation Sites that diminish at 48 hpi. **a**, 9 hpi, wild type (WT). **b**, 9 hpi, integrase mutant (IN). **c**, 48 hpi, wild type. **d**, 48 hpi, integrase mutant. All plots are generated with a k-mer length of 27 and a base offset of 455 (see Supplementary Material) to match the standard positions of the HIV genome. *Coverage ratio is plotted with Bayesian regression and log transformation. A prominent terminus signal observed at 635 on the negative strand in this analysis corresponds to the previously well-described initiation site for HIV minus strand at the tRNA primer binding site, while the prominent terminus at 9074 on the plus-strand corresponds to the previously well-described terminus reflecting the left end of the U3 segment of the LTR.

Our analysis revealed a number of additional terminus signals of lower magnitude but with consistency across replicates. One such signal, consisting of a significant end capture signal at base 4800 and a subsequent strand bias peak just downstream, corresponds to an additional feature that has been studied during HIV plus-strand synthesis: the central polypurine tract, or cPPT.^36^ Consistent with the previously proposed flap architecture at the cPPT^36,46^, sharp peaks were observed for positive strand bias and positive start site enrichment at position 4800 corresponding to the cPPT. In contrast, signals for both strand bias and start site enrichment on the negative strand were more distributed. This is consistent with the lack of a defined terminus on the negative strand per the flap architecture.

The flap has been proposed to play several roles in HIV replication, including in nuclear import and/or initiation of plus-strand synthesis.^43,47–49^ We further note the substantial decrease in both end capture and strand bias signals at 48 hpi compared to 9 hpi (Figs. 4c-d). This is consistent with a reduction in the extrachromosomal linear dsDNA form at the 48 hpi time point compared with the 9 hpi time point, which is expected based on the known kinetics of HIV genome replication.^50,51^ The sequence upstream of the strong terminus at 4800 is shown in Table 1, with a polypurine tract immediately preceding the prominent terminus. This configuration suggests that the proposed polypurine RNA primer left after Ribonuclease H (RNAse H) activity of HIV reverse transcriptase has been trimmed in vivo and/or during repair prior to sequence capture in generating the next-generation sequencing libraries used in these analyses, so that the eventual terminus in the library is representative of a discontinuity between RNA and DNA sequence during plus-strand synthesis.

Of substantial interest, the site at position 4800 is not the only point in HIV where a distinct end capture signal is followed by some *k*-mer strand bias signal and where these are decreased at the 48-hour time point following resolution of intermediate structures formed during the early infection process. Additional sites with similar characteristics are observed at several sites to the left and right of 4800 bp, suggesting that there may be additional sites with similar roles and characteristics in HIV plus-strand synthesis. Beyond the signals for the LTR termini and the central flap, the next-most prominent signals were present at positions 6266, 1137, 5279, 7592, and 8428, with several additional sites observed throughout HIV (Table 1). These five most prominent sites are all termini facing in the positive direction, and notably all are at the end of purine-rich stretches. A weaker pair of terminus sites on the minus-facing direction (6339/6340, Table 1), adjacent to the 6266/6267 plus-facing termini, appears further down in the list (as ranked by capture frequency).

Given the preponderance of terminus signals present on the plus-strand, preceded by a polypurine tract, and depleted in data from the 48-hour sample, we hypothesize that these elements could play a parallel role to the cPPT. We provisionally denote these positions as altPPTs (alternative polypurine tracts) because they were generally located in purine-rich regions (Table 1). The top five altPPTs are unusually purine-rich (75% - 93.75%). A calculated chance of five arbitrary plus-strand 16mers derived from HIV having >75% purine content) is approximately 1 in 10,000 (see Supplementary Material).

One possible biological implication of the altPPTs is that they might serve as additional sites for initiation of plus-strand DNA synthesis during HIV replication (Fig. 4, Table 1, and Fig. S1). We note that the observed peaks are highly reproducible in replicate datasets, and that the 9 hpi and 48 hpi samples were prepared in an identical manner for sequencing. These observations argue against technical artifacts such as sequence-dependent DNA fragmentation biases, as these would be insufficient to explain why the signal for altPPTs is differential between the 9 hpi and 48 hpi time points (Figs. 4c-d and Table 1). Both integrase mutant and wild type samples exhibit differential enrichment for the altPPTs at the 9 hpi time point compared to the 48 hpi time point, although there is generally slightly lower start site enrichment in integrase mutant samples compared to wild type (Fig. 4 and Table 1). This analysis highlights the potential of end detection to define subtle differences in DNA structures that may illuminate biological function.

## Discussion

Understanding the configuration of DNA ends in a biological sample is a key component in characterizing genome structure and dynamics in the underlying biological material. While direct end capture methods exist, there are cases where capturing the end directly can be difficult. In this work, we have added an indirect means to detect DNA termini, using the asymmetry in calculated coverage resulting from the intrinsic read and fragment lengths in high throughput sequencing experiments. This metric, strand bias, has previously been discarded in some cases as an artifact arising from systematic errors and used to discard “low-quality” samples.^21,52^ On the other hand, logic analogous to strand bias has been observed to correspond to the starts and ends of transcripts in RNA sequencing data and in other special cases.^22,23^ Given several means to detect ends, it is valuable to observe both the end capture and strand bias signals simultaneously. We applied such analysis to a number of high throughput sequencing datasets, with results illuminating both opportunities and limitations of the combined approach.

We first evaluated bacteriophage lambda to determine if a well-defined *in vitro* sample could be analyzed this way. We were able to readily detect ends, including both the genomic DNA ends as well as the restriction enzyme-generated cut sites. Our exploration demonstrated that the ability to subtract a control sample, corresponding to the uncut DNA, helps minimize noise and get clearer signal from genuine ends.

Next, we sought to establish whether our method could reliably detect the ends of an *in vivo* sample with a blocked end, in the case of bovine adenovirus. As expected, we did not observe capture of the ends, but we did observe the known ends of the adenovirus as strand bias peaks, demonstrating the value in combining both approaches.

### Application to HIV replication

To examine a more complex condition and see if we could gain novel insights, we further applied our methodology to HIV. In addition to identifying the known 3’ and 5’ ends of the linear unintegrated dsDNA form of HIV in cells, we also observed other regions of prominent termini and strand bias that may potentially play roles in HIV replication. Particularly interesting in this context were a set of purine-rich sequences (PPTs) on the sense strand, including both a known central site (cPPT) and an additional series of similar secondary sites, which we refer to as altPPTs. Our data suggest that the role of the cPPT may not be played by just a single sequence but potentially by several sequence elements.

Although its precise role during early stages of HIV replication is debated, the cPPT is required for optimal viral replication^53,54^ and its presence in lentiviral vectors ensures high transduction efficiency^55-58^. As PPTs are purine-rich, they resist digestion by RNAse H, allowing them to persist and serve as primers for the next step in replication.^59^ They can serve as an entry site for initiating plus-strand synthesis, allowing protection against host defenses that target single-stranded DNA by accelerating the transformation to double-strandedness of the full genome (Fig. 5).^43,48,60^

**Figure 5.**
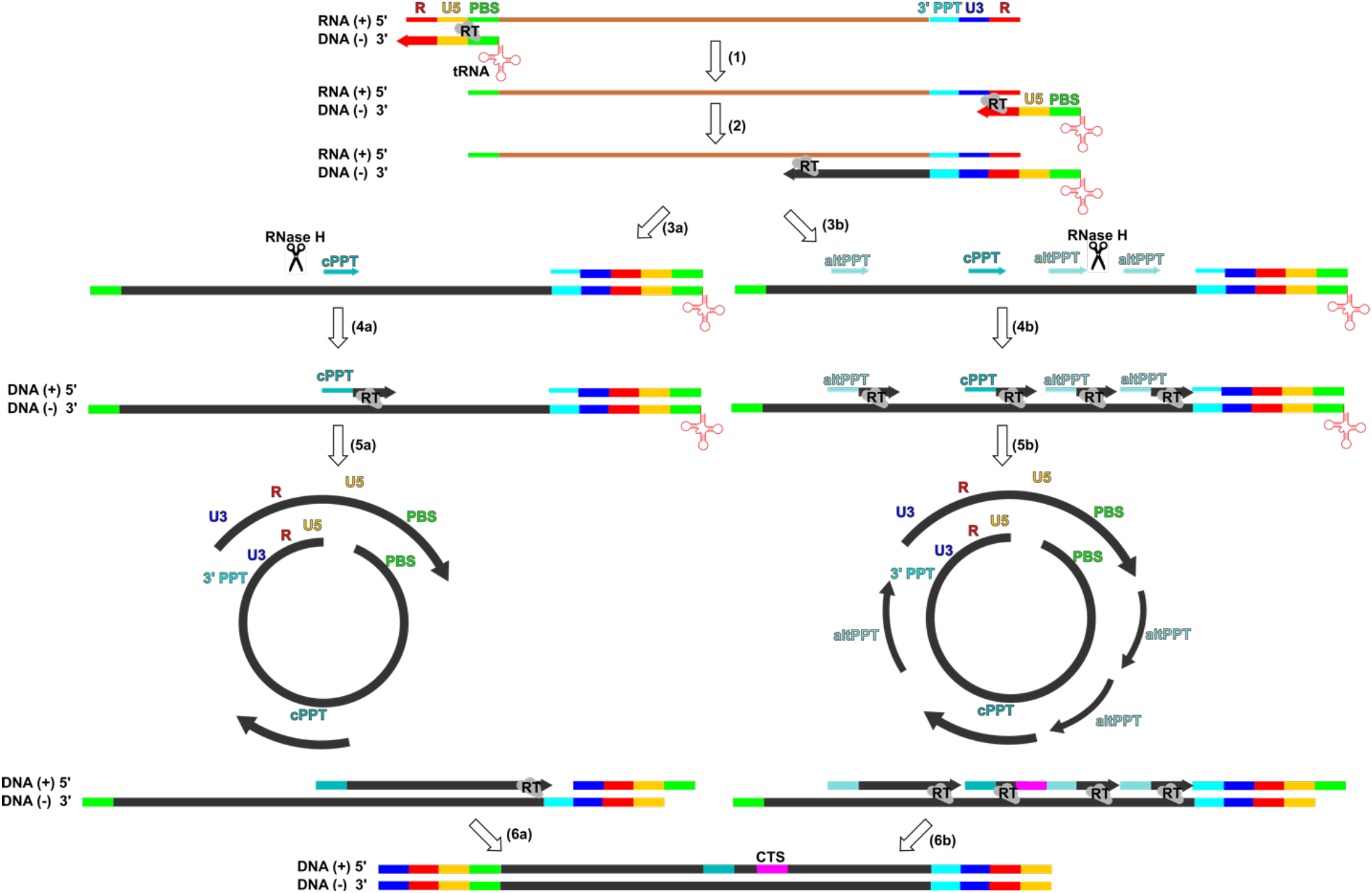
Current and Proposed Mechanisms for HIV Replication and potential roles for alternative PPT sites (altPPTs) in HIV plus-strand synthesis. 1) tRNA primer at the 5’ PBS site initiates reverse transcription. 2) First jump: R’U5’ is moved to the 3’ end of the genome and triggers minus-DNA synthesis. 3a-5a) Current lentiviral replication mechanism: 3a) RNAse H cleaves all the RNA except for the cPPT and 3’ PPT, 4a) Both the cPPT and 3’ PPT are used primers to initiate plus-strand DNA synthesis, 5a) Downstream plus-strand DNA is synthesized until the RTase reaches a strong-stop DNA site (U3-R-U5). 3b-5b) Proposed HIV replication mechanism: 3b) AltPPT sites are protected from RNAse H cleavage, in addition to the cPPT and 3’ PPT, 4b) Multiple altPPTs can serve as initiation sites, in addition to the cPPT and 3’ PPT, for plus-strand synthesis, 5b) With multiple initiation sites (altPPTs), synthesis of downstream plus-strand DNA is accelerated. 6) Ultimately, the synthesis of upstream plus-strand DNA stops at the CTS site near the genome’s center, creating a discontinued plus-strand DNA.

The proposal of multiple plus strand initiation sites in HIV is consistent with an earlier multisite initiation model derived through a kinetic analysis of reverse transcription by Thomas *et al*.,^61^ but to our knowledge the relevant sites in HIV had not been previously defined.

Treatment with the HIV replication inhibitor SJP-L-5 has been shown to correlate with appearance of multiple fragments proposed to reflect multiple initiation sites for plus-strand DNA synthesis.^60^ It is conceivable that the multiple sequences we observe here (altPPTs) reflect the additional initiation sites for plus-strand synthesis where signals are enhanced following SJP-L-5 treatment (Fig. 5, Steps 3b-5b); but we currently have no evidence to support any connection.

Multiple initiation sites could certainly contribute to the overall speed of the reverse transcription process. Beyond the overall acceleration, becoming double-stranded sooner would protect HIV from single-stranded DNA editing by APOBEC3 proteins.^43,48^ AltPPTs may also exist in other retroviruses, as multiple single-stranded regions per linear viral DNA have been observed in Rous sarcoma virus.^62^

### Limitations and caveats

Given that library protocols are (highly) varied and that many interesting genomes have inherently complex and repetitive sequences, particularly at the ends, strand bias and end capture may lead to different results with different datasets. There are structural features that are not simple termini that would affect the ability of sequencing and capture in those regions both positively and negatively, potentially yielding false terminus signals. In particular, repetitive regions may lead to artificial biases reflecting issues in the alignment process, and therefore must be analyzed with caution.

Providing a set of real-world challenges, mitochondrial DNAs exist in both linear and circular forms, with topological distribution differing across species, and with many mitochondrial genomes having sequence irregularities that would complicate sequencing-based terminus identification.^63^ Taking two standard models as examples, there are long terminal inverted repeats at the ends of the linear *Chlamydomonas reinhardtii* mitochondrial genome,^64,65^ while the circular *Caenorhabditis elegans* mitochondrial genome has complex repeats in an AT-rich region of the circle.^66^ Although circular, *C. elegans* mitochondrial DNA may also replicate via a rolling circle mechanism that produces linear concatemer products.^67^ Illustrating the challenges of templates with complex structure (both in base composition and repetitive character), we provide an exemplary description of some of these challenges in the supplementary material (Figs. S2-S5).

While the combination terminus-detection and strand-bias approach could detect any form of linear DNA in principle, it is particularly suited for linear extrachromosomal DNA, because the sequence ends are typically well-defined. In contrast, linear chromosomal DNAs generally have terminal telomere structures, which exhibit length variability in the terminal tandem repeat^68,69^, presenting a challenge for sequence alignment. The method is also currently not applicable for much larger whole chromosomes, such as those in humans, as the complexity of these chromosomes greatly increases the likelihood of confounding sequence-dependent effects on capture and sequencing. More efficient experimental enrichments, alignment algorithms and/or strategies that only examine smaller regions of DNA may help facilitate work in this area.

Finally, while control conditions are helpful to amplify signal, such as in the case of lambda (uncut versus cut) and HIV (earliest versus later stages in reverse transcription), such controls are not available in all published datasets, and care must be taken to distinguish genuine peaks resulting from DNA termini versus peaks that arise from fragmentation biases.

### Conceptual considerations in inference of ends from next-generation sequencing of DNA fragment pools using end capture

Sites with greater-than-expected end capture are candidates for true ends in DNA. In addition to such *bona fide* ends, ends are also formed by fragmentation, but any kind of reference sample or model for expected end incidence can be checked against real data. While other processes can result in preferential end capture, such as greater DNA fragility, better capture, or more effective amplification or sequencing, these sites nonetheless serve as candidates for possible ends.

While direct end capture is useful in many scenarios, preferential end capture can only happen with certain sequencing approaches. In particular, libraries prepared with Nextera, which uses enzyme-based tagmentation to fragment DNA, would have less distinctive end capture signals.^70,71^ Despite this, methods like Nextera may still result in strand bias near the ends, leading to a potentially useful readout of termini.

## Conclusion

To detect DNA termini and linear extrachromosomal DNA from next-generation sequencing of fragmented and captured dsDNA, we proposed and tested a method based on combining a direct approach assessing preferential end capture and a novel indirect approach using strand bias.

Termini are reflected by high start site enrichment and a strong strand bias, with inward-facing reads greatly enriched over outward-facing reads in the immediate proximity of any end. This technique can be employed to nominate linear extrachromosomal DNA for a wide variety of applications, and there are numerous scenarios where such analyses would be useful towards identifying ends and linear extrachromosomal DNA, including cases in which a reference genome is known or unknown or investigating conditions in which ends may be generated unexpectedly.

Applying this method to HIV identifies known termini and confirms a previously described unique structure, the HIV flap sequence. More importantly, we have discovered multiple purine-rich sequences (polypurine tracts – PPTs) on the plus-strand, which we call alternative PPTs (altPPTs), as putative initiation sites during plus-strand synthesis that might contribute to HIV replication.

## Methods

### Lambda DNA digestions and sequencing

Lambda DNA (NEB cat # N3011S) was digested for 3 hours with PvuII-HF (NEB cat #R3151S) or PstI (NEB cat #R0140T) in 50 ul reactions. 450 ul of TE pH 8.0 was added and digested DNA was extracted with equal volumes of phenol:chloroform and subsequently chloroform. 40 ul of saturated ammonium acetate was added to aqueous phase prior to precipitation with 100% ethanol. The DNA pellet was washed with 100% ethanol and resuspended in 25ul of TE pH 8.0. Sequencing libraries were prepared using Nextera XT (Illumina cat # FC-131-1024) and 200 bp-500 bp fragments were cut from a 1% agarose gel. DNA sequencing was performed on an Illumina Miseq instrument. Data will be available at the Sequence Read Archive.^72^

### Bioinformatic analysis

Datasets were first downloaded from the Sequence Read Archive^72^ and then analyzed in parallel using two distinct software pipelines.

For alignment-based approaches (exemplified in Figure 2), adapter trimming using *Trimmomatic* (version 0.39) ^73^ was followed by alignment to the reference sequence using *Bowtie 2* or *BWA*, producing a SAM file.^25-27^ Next, we used *SAMtools* to convert the SAM file to a BAM file, which was sorted and indexed, then analyzed via *Pysam*.^74^ From here, coverage across the genome was used to compute strand bias, which is plotted, in some cases, applying filtering for positions where the total number of reads meets a specific count threshold (not applied for Fig 2 in this paper but useful for datasets and regions with limited coverage).

*PolyBench* provides a single-step kmer based approach that takes raw sequence reads and a reference file as input and yields graphic and tabular output on strand bias as well as a number of other positional and global run characteristics. *PolyBench* is designed for relatively short reference sequences (generally <100kb) and can generally be run on a personal computer or lab workstation (standard settings were klen=27 StartEndPlot=True). *PolyBench* uses a simple Bayesian regression tool (*BayesRatio*) to handle low coverage areas of the genome, where strand bias is not as informative due to stochastic variations that preclude a stable +/-ratio (*BayesRatio* estimates the power of the data from discrete measurements under two distinct conditions to distinguish an arbitrary fold-difference; *PolyBench* uses this calculation to regress plotted values to the minimum-fold difference that would be consistent with a provided false discovery ratio.

Separate code for this calculation is available at GitHub/FireLabSoftware/BayesRatio).

### Considerations for the use of strand bias as a tool in detecting termini

The differential plus and minus strand bias reflects a complex aggregate value at each *k*-mer or position--how often a fragment is captured and a read obtained that covers that position (which we call position *x*). The requirements for this are

1. In addition to the end at *x*, the “other end of the fragment” must be sufficiently far away to allow a captured fragment to fit within the distance between the two. A fragment that is too short to be captured using the sequencing methodology in use will be missed by this analysis.
2. The full intervening sequence between these two ends needs to be “clear”, “intact”, and “sequenceable” to allow amplification and sequencing (i.e., minimal fragmentation breaks or roadblocks)

Note that in addition to the end captured at *x*, another captured end may similarly be present at the same point and facing the other direction, and that this would provide a separate (and somewhat confirmatory) signal if present.

If capture and sequencing are uniformly insufficient, the data may be noisy, but there could still be sufficient evidence to detect differential representation, particularly if there are uncleaved control samples, expectations of where termini could occur, or multiple samples for confirmation. For DNAs and methodologies where the resulting data are biased, however, there may be substantial challenges in detection of *bona fide* termini. In particular, if the relative amounts of DNA, rates of end capture, and recovery of sequences are positionally determined reflecting DNA structure or sequence, there can be differential plus and minus effects at a given position. This is conceptually determined by the fact that the plus-strand *k*-mer or alignment count represents the (presence + capture + recovery + sequencing) of fragments that start just to the left of *x* and extend rightward, while the minus strand values represent the (presence + capture + recovery + sequencing) of fragments that start just to the right and extend leftward.

A few influences on this are of note:

1. DNA abundance can be affected by differential presence of the DNA (i.e., there can be real differences in DNA abundance in a sample because of amplification, deletion, aneuploidy, etc.)
2. End capture can be made non-uniform by pre-existing ends (leading in some cases to a greatly increased capture rate at a single site (if the resulting end can be captured) and in others to a decreased rate of capture relative to unbroken DNA (e.g., for methodologies that require a dsDNA region on both sides to insert a tag for ends that are blocked).
3. Regions that resist the fragmentation methodology used in the library construction may influence the distribution of cuts that can generate either the proximal or distal ends for sequencing. Depending on the degree of fragmentation and the length distribution of recovered fragments, highly susceptible regions can either increase or limit the recovery of appropriate length fragments for sequencing (similar for regions with less-than-average susceptibility). Modifications to the DNA (or damage) can also affect this, and sequence-biased damage, modification, or sequence features that are recalcitrant to sequencing or amplification (homopolymeric runs, hairpins, etc) would have a significant effect here.
4. Library construction tools impose different degrees of length constraint on the eventual sequenced fragments. For methodologies that impose only a broad requirement on fragment length, the effects of the plus-downstream or minus-upstream sequence cut requirement may be limited during averaging out over a longer region. For methods that impose a precise requirement on fragment length, the other-side-of-the-fragment could substantially influence relative recoveries.
5. Some sequences can impede or interfere with sequencing reactions and if these are strand specific or polar (e.g., on one side of position *x*), there would certainly be consequences for the differential plus/minus ratio.

For a DNA sample and library/sequencing methodology where region/site specific effects on sequencing depth are minimal (i.e., ends in the DNA are the only features of the above with a substantial region-specific effect), the strand bias (plus coverage/minus coverage) should perform extremely well. However, this will never be perfect; in some cases, the resultant noise is tolerable in detecting the large signal resulting from breaks. In other cases, we will have an “uncut” (or unfragmented) DNA as a control for sequence-specific effects and a likely ability to subtract such effects. But for datasets where there are substantial positional effects on capture and sequencing, we might expect some specific features to the noise.

### Offset signals in strand preference analysis

One feature to be expected comes from the offset between the area relevant to plus-strand sequence recovery and the area relevant to minus-strand sequence recovery.

Plus-strand recovery for a *k*-mer at position *x* reflects characteristics of a segment that would be of length *FragmentLength* and start either right at *x* or slightly upstream (so that *k*-mer *x* would still remain within the read). Numerically this sets the left edge of fragments that can result in a “plus” count of the *k*-mer between (*x* – *ReadLength + KmerLength*) and position *x*.

Minus strand recovery reflects characteristics of a segment that will be to the right of this. A similar calculation has the left end of this segment between (*x – FragmentLength + KmerLength*) and (*x* – *FragmentLength + ReadLength*).

The offset between these two ranges is (*FragmentLength* – *ReadLength*). This could be non-evident if either the fragment length is quite variable or the underlying sequence representation methodology is quite uniform (the former would diffuse the offset signal out over a large set of offsets, and the latter results in less of a signal to be observed). But for experiments where a narrow range of fragment sizes is selected and there is quite a bit of underlying nonhomogeneity in representation along the sequence, we will see the kind of offset pattern observed with the *C. elegans* mitochondrial sequence dataset shown in Supplementary Material (Fig. S3). There will certainly be other, more local, effects on the signal of plus/minus (true ends in the DNA but also micro-heterogeneity in capture/sequencing, DNA damage, underlying sequenceability, etc), as well as stochastic effects. Therefore, the offset may be subsumed by background in many datasets.

The offset noted in the comparison in Fig. S3 was 413 bp, the read length in the sample was 101, and Goldstein *et al*.^75,76^ noted in their description that they isolated fragments with a target size of 501 bp. For most separation methodologies, the difference between 500 and 514 as-good-as-can-be-expected. Thus, the offset may be the juxtaposition of an unusually precise sequence-length selection and an unusually-nonuniform-capture methodology, which may be lurking in all datasets to some extent.

## Data availability

Published sequence datasets described herein for bovine adenovirus, *C. reinhardtii, C. elegans*, and HIV were obtained from the NCBI short read archive with accession IDs DRR257739 (bovine adenovirus 7^34^) and SRR879372 (*C. reinhardtii)*, SRR7211661/SRR3536210/SRR7211661 (*C. elegans*^*75,76,79,80*^), and PRJNA559271 (HIV 9 hpi WT, 48 hpi WT, and 9 hpi IN). Additional HIV datasets and sequencing data for full length and cleaved bacteriophage lambda are being provided to NCBI. Reference genomes for bovine adenovirus, *C. elegans*, and *C. reinhardtii* were from GenBank at, LC597488.1, NC_001328.1, U03843.1 respectively.

## Code availability

The programs used to generate the figures in this paper are available on GitHub at https://github.com/FireLabSoftware/.

## Acknowledgments

We would like to thank Virginia Walbot for comments on the manuscript, and members of the Fire Lab (Orkan Ilbay, Ivan Zheludev, Emily Greenwald, Usman Enam, Dae Eun Jeong, Janie Kim, Matt McCoy, Massa Shoura, Nelson Hall, Amisha Kumar, Sameer Sundrani, Lorenzo Angcanan Del Rosario, Lamia Wahba, and Loren Hansen) for their help and support. This work was supported by the National Institutes of Health (NIH Award no. R35GM130366) and the Stanford University Program for Bioengineering, Biosciences and Biomedicine (Bio-X). The work in M.B.’s lab was supported by grants from European Research Council ERC-2018-ADG (RetroChrom#835184), MSD-Avenir, GILEAD, Labex EpiGenMed “Investissements d’avenir” (ANR-10-LABX-12-01), and Fondation pour la Recherche Médicale (FRM; LEQ20151134476). S.M. was supported by National Center for Global Health & Medicine Research Institute, JSPS Overseas Research Fellowships (Japan Society for the Promotion of Science), Mochida Memorial Foundation, Uehara Memorial Foundation, Kato Memorial Bioscience Foundation, Osamu Hayaishi Memorial Scholarship for Study Abroad, Yamada Science Foundation and ERC-2018-ADG (RetroChrom#835184).

## Supplementary Material

**Fig. S1.**
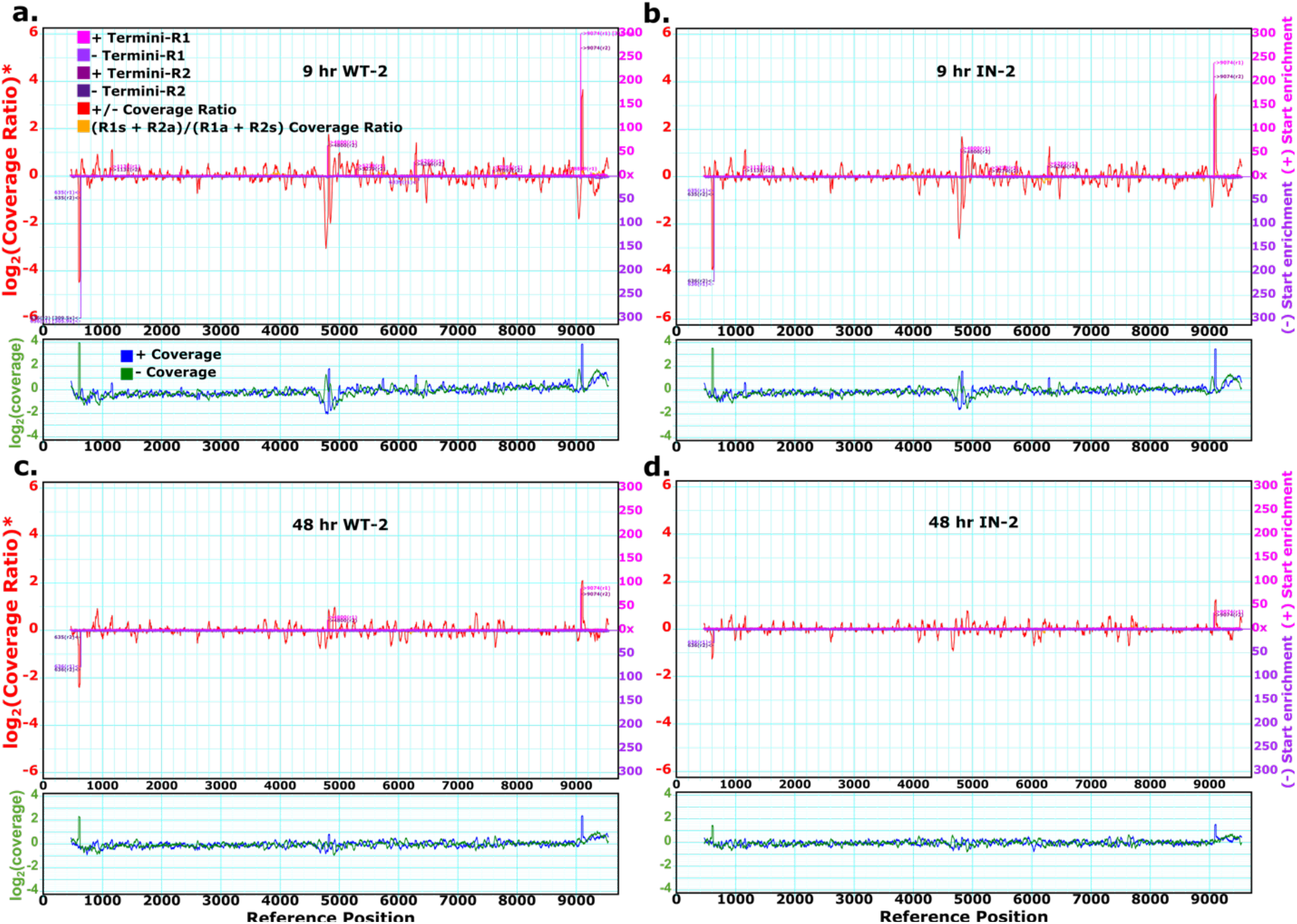
Second replicate: Strand bias and end capture of the HIV genome, calculated by PolyBench, demonstrate DNA termini and [Putative] Plus-Strand Initiation Sites that diminish at 48 hpi. **a**, 9 hpi, wild type (WT). **b**, 9 hpi, integrase mutant (IN). **c**, 48 hpi, wild type. **d**, 48 hpi, integrase mutant. All plots are generated with a *k*-mer length of 27 and a base offset of 455 to match the standard positions of the HIV genome.

### Assembly of a canonical HIV genome copy based on viral products derived from clone pNL4-3

The template for the production of the virus used in the studies of Machida *et al*.^35^ was the reconstructed full-length HIV genomic insertion clone pNL4-3 of Adachi et al.^77^ Because the upstream and downstream LTRs in pNL4-3 are distinct in sequence, a careful reconstruction of the expected sequence in LTR regions was needed as a basis for expectations of derived DNA sequences following virion infection. To obtain such sequence, MEGAHIT assembly of sequences from the HIV datasets was followed by appropriate permutation of the termini to produce an appropriate reconstruction of the starting RNA template for reverse transcription.^44^ Because the initial RNA transcript starts within the first LTR at position 456,^78^ we permute the resulting assembly as expected for the known boundaries of terminal duplication for HIV. The resulting model for linear dsDNA consists of three segments (i) a segment spanning base 1 to 455 of the right LTR from pNL4.3, (ii) a segment spanning base 456 of the left LTR through base 455 of the right LTR, and (iii) a segment spanning base 456 of the left LTR through the end of the left LTR. This sequence has been denoted pNL43_Transcript.fa. Note that analysis of the sequencing data also allowed identification of the inactivating mutation in the INtegrase mutant datasets as D116N.

### Probability of purine richness of altPPT Sites

The probability that any given 16-mer in the HIV reference genome we assembled is at least 75% purine-rich and ends in a purine site is approximately 15.86% (see the attached program, hivsim.py). 0.1586^5^ ≈ 1.003 × 10^−4^, which is around 1 in 10,000. Since the purine richness for four of five altPPTs is greater than 75%, the true chance will be much lower than this.

### An example of methodological complexities introduced in experiments with repetitive genomes and nonuniform capture

We describe analyses of three datasets consisting of *C. elegans* DNA which exemplify complexities that can arise from nonuniform capture and genomic areas with unusual sequence characteristics (Figs. S2-S4). We start with the caveat that while the *C. elegans* mitochondrial genome is known to be circular under some conditions, there may be times when there are linear molecules; indeed, it has been shown that *C. elegans* mitochondria can utilize a rolling circle mechanism resulting in linear concatemers.^67^ From these three (rather arbitrarily chosen) datasets obtained for *C. elegans* genomic sequence, we found rather distinct patterns as a likely result of technical differences and variation, which are illustrative of caveats in interpretation with different datasets. The first *C. elegans* sample (Fig S2) provided relatively constant coverage throughout the genome with modest strand biases at several sites, and with a dramatic drop in coverage toward the right of the standard mitochondrial map. While the differential strand coverage, particularly combined with specific end capture, can evidence a distinctive end, datasets where there is substantial variability (and many prominent ends on both strands, with no control of presumably-intact DNA under the same conditions) leave questions about whether there are *bona fide* termini for each sequence anomaly. The far-right end is particularly notable here. This highly AT-rich region produced a lower signal, potentially reflecting a mixture of capture, sequencing, and positioning challenges, and making this a particularly difficult region to interpret signals. A second dataset from the same organism using a different capture method (in this case ligation based) showed an even more dramatic set of differential stand bias peaks (Fig. S3). As noted in the Methods section, the strand bias here appears consistent with a combination of highly uniform fragment sizes and highly variable yield across regions of the genome; with no strong reason to suspect that these are *bona fide* ends. A third *C. elegans* dataset (Fig S4) shows rather uniform coverage with no strong terminus signals in the majority of the sequence and just the anomalies in the AT-rich region shown at the right on the plot. Although the distinctions between *bona fide* termini and other phenomena eventually will rely on additional methods of determination, we suggest that uniformity of coverage in regions where no terminus is expected can serve as a preliminary indicator of whether bias signals around potential termini would merit further investigation.

**Fig. S2.**
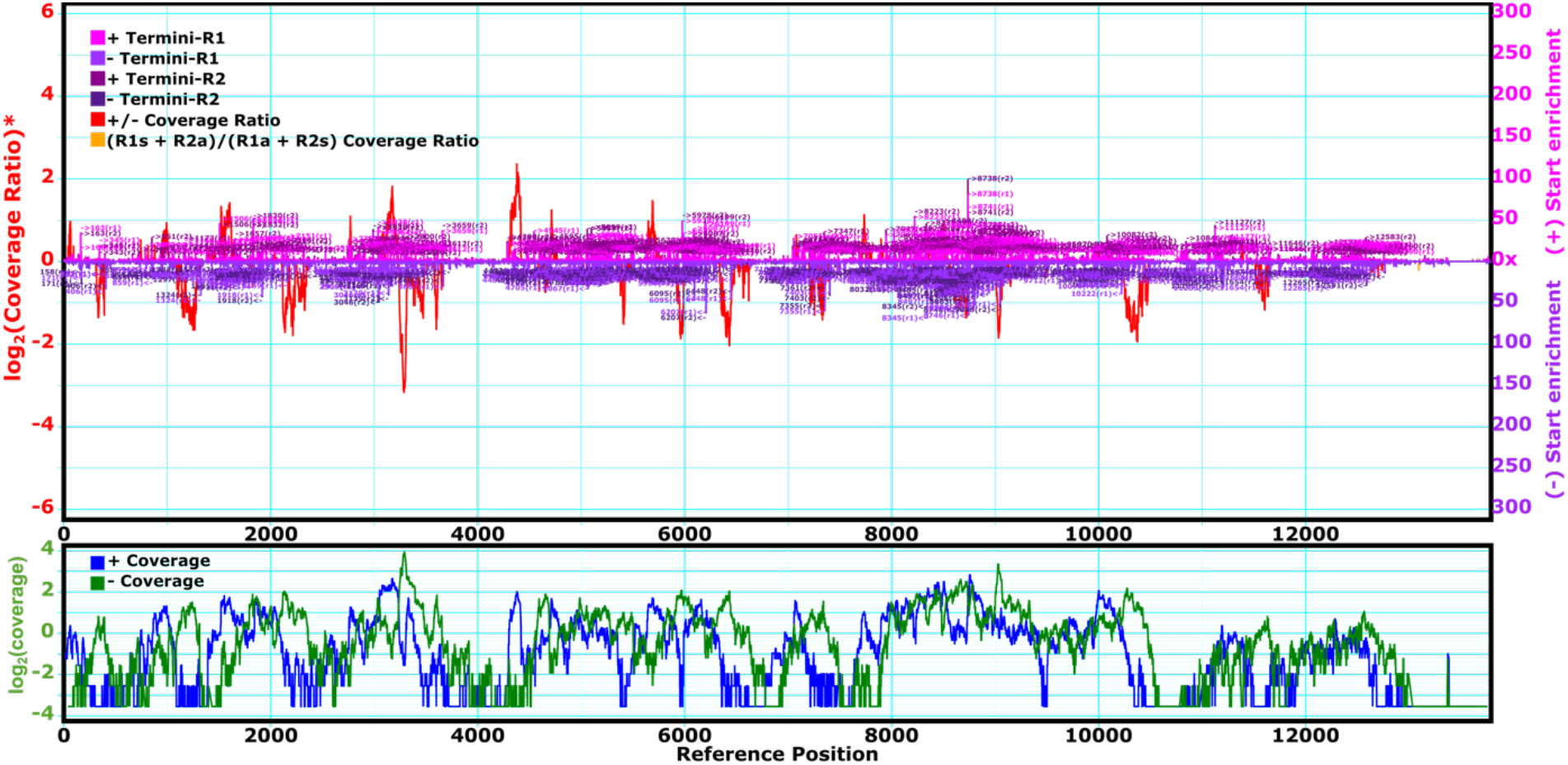
Strand bias and end capture of a *C. elegans* mitochondrial sequencing experiment. This particular dataset (SRR7211661)^79^ was obtained with a library prepared with Nextera tagmentation. Coverage is nonuniform across the sample and numerous peaks are observed in both end capture and strand bias profiles. The overall nonuniformity in coverage would be consistent with numerous ends being present, the data would also be consistent with non-termini-dependent biases due to nonuniform capture.

**Fig. S3.**
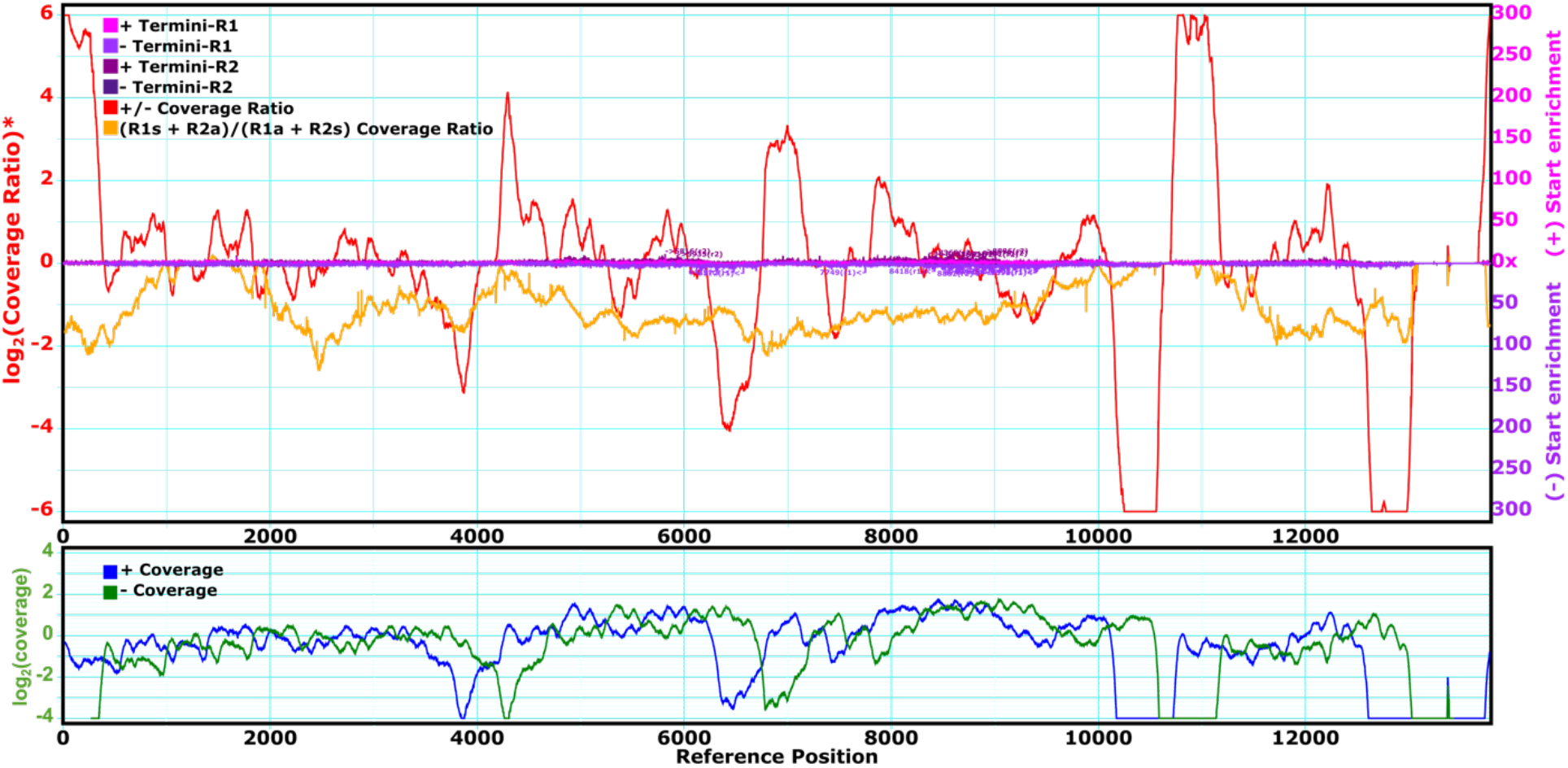
Strand bias and end capture of a second *C. elegans* mitochondrial sequencing experiment. The dataset (SRR3536210)^75,76^ was obtained using a library prepared using DNA fragmentation and careful size selection. The precise size selection combined with severe drops in coverage at several points leads to an offset between sense and antisense coverage and hence to dramatic peaks in the strand bias plot, which would seem unlikely to correspond to true ends. Figure S5 shows an additional example, a case of a characterized linear genome (*C. reinhardtii* mitochondrial genome). In the case of this dataset, the local coverage is uniform for much of the genome, suggesting that termini might indeed be evident in further analysis of the underlying sequence data. However, the presence of inverted terminal repeats on the linear molecule leads to complexities in counting coverage in the relevant regions. Anomalous strand ratios are observable in the terminal repeat regions, but the conflation of *k*-mers from left and right end repeats (due to the inverted nature of the repeats) makes a clear definition of the linear structure difficult.

**Fig. S4.**
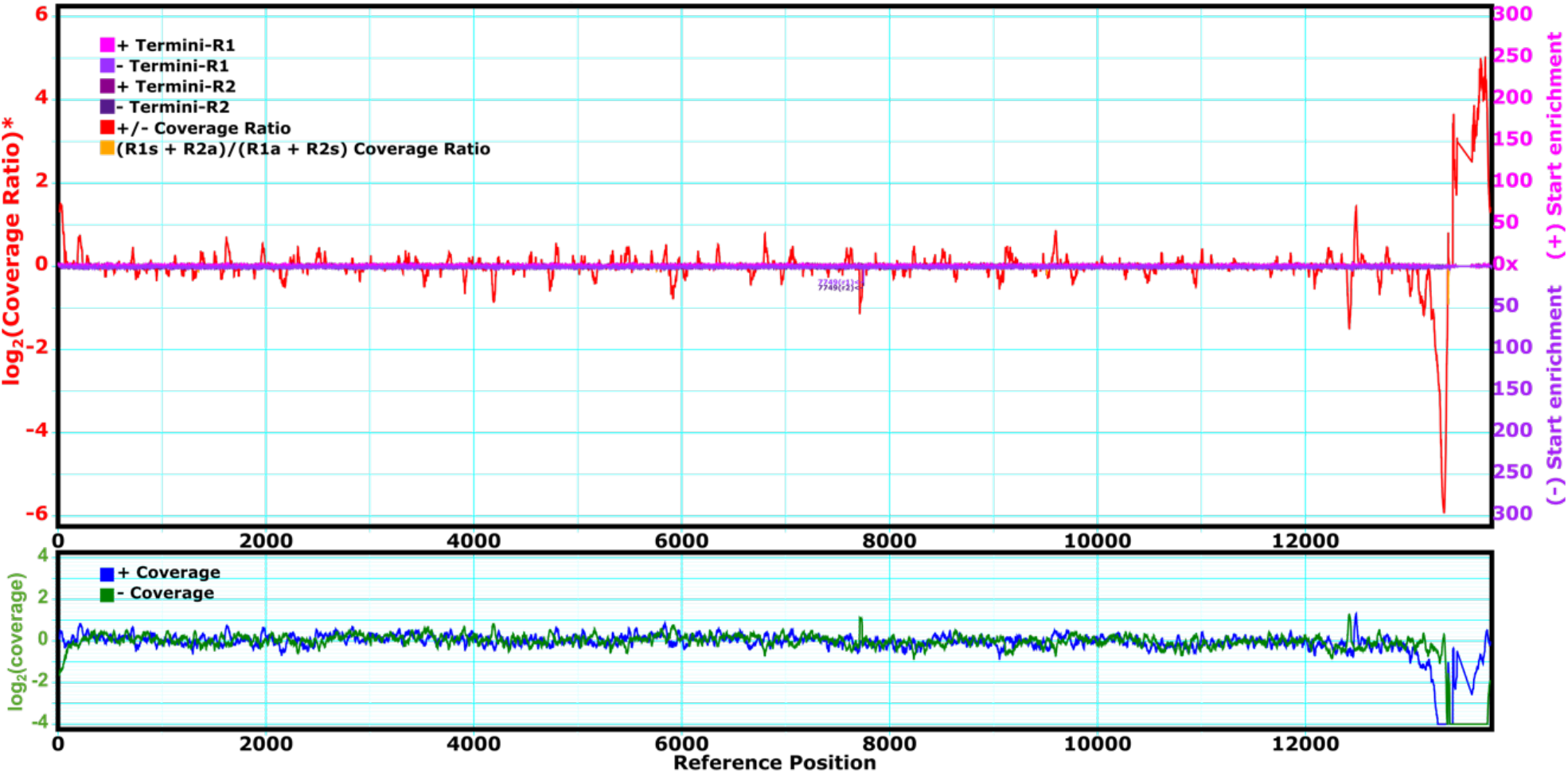
Strand bias and end capture of a third *C. elegans* mitochondrial sequencing experiment. The dataset (SRR8176698)^80^ was obtained using a library prepared using TruSeq. While there is little signal outside from scattered noise for most of the genome, the far-right AT-rich end presents difficulties in alignment leading to specious peaks.

**Fig. S5.**
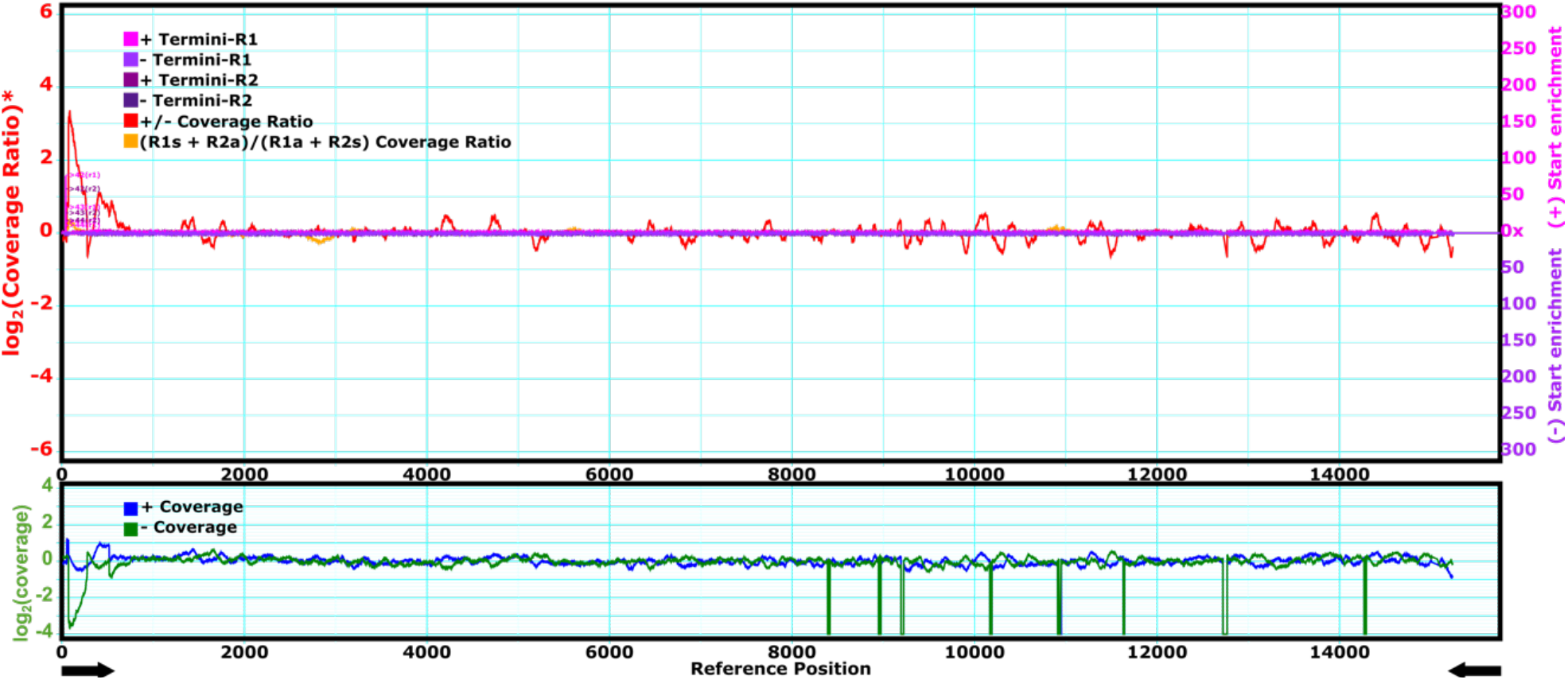
Strand bias and end capture for mitochondrial DNA from a *C. reinhardtii* sequencing experiment prepared using Takara PrepX, calculated by PolyBench. Minimal signal is displayed except at the left end where a terminal repeat is present. Points where coverage dips to zero here may represent variation between the published reference sequence and the underlying sequence of the cultivar used for these experiments, while the strand bias peak being limited to the left end may be the result of an inverted duplication at the ends of the sequence (all *k*-mers from either end would map to the left side here as a result of Polybench’s first-match handling of repeated sequence in a reference). Arrows indicate the inverted terminal repeats at the ends. Data from Perlaza et al^81^.

